## Supplementary material for "DREAMS: Deep Read-level Error Model for Sequencing data applied to low-frequency variant calling and circulating tumor DNA detection"

### Supplementary section 1: Sample collection and preparation

**1.1 Sample collection and processing**

Patient tumor and blood samples were collected as part of the IMPROVE studies (NCT03748680) at Aarhus University Hospital, Denmark. Tumor samples were collected as either formalin-fixed paraffin embedded (FFPE) samples or fresh frozen tissue. Pre- and post-operative blood samples were collected in EDTA tubes and fractionated into plasma and buffy coat within 2 hours of collection by double centrifugation for ten minutes at 3000*g*.

DNA from FFPE and fresh frozen tissue was extracted using the QiAamp DNA FFPE tissue kit (Qiagen) or the Puregene DNA purificaition kit (Gentra Systems), respectively. Germline DNA was purified from buffy coat using the QiaAMP DNa Blood Medi kit (Qiagen). DNA from tumor and buffy coat was quantified using Qubit (Thermofischer), and stored at -80°C.

Cell-free DNA (cfDNA) was extracted from 8 mL of plasma using the QIAsymphony DSP Circulating DNA kit for the QIAsymphony instrument (Qiagen). cfDNA was quantified using ddPCR as previously described [1].

**1.2 Library preparation**

Prior to library preparation, DNA from tumor and buffy coat was fragmented by sonication (Covaris E330) to a target size of 300-400bp. Library preparation was performed according to manufactures protocol using the KAPA Hyper Library Preparation Kit (KAPA Biosystems) with minor adjustments as previously described [2]. Adjustments included; 1) increased ligation time of 30 min at room temperature, 2) increased DNA:beads ratio to 1.4x after adapter ligation to ensure retainment of small fragments and 3) PCR amplification of libraries was split into four reaction to limit PCR bias. Final libraries were quantified using Qubit (Thermo Fisher Scientific).

**1.3 Targeted sequencing**

To facilitate high depth sequencing, libraries were enriched for regions of interest using a customly designed capture panel. The capture panel (SeqCap EZ Enrichment, Nimblegen) was designed to include the most commonly mutated regions in colorectal cancer based on TCGA data[3]. Targeted regions included large parts of TP53 and KRAS along with frequently mutated sections of SMAD2 and SMAD4, APC, BRAF and PIK3CA among others (Supplementary Table 1). Additionally, 18 single-nucleotide-polymorphisms (SNPs) positions were included in the panel to ensure that tumor, buffycoat and plasma sample sets originated from the same patient (Supplementary Table 1).

Capture of target regions was performed following standard SeqCap EZ HyperCap protocol by mixing 2-8 libraries in each capture reaction. The entire capture protocol was performed twice using 0.5x probe for each capture. This ensured a high fraction of on-target fragments without increasing the amount of used probes. Captured samples was quantified using TapeStation HS (Agilent) and subsequently paired-end sequenced (2x150 bp) on NextSeq 550 (Illumina).

### Supplementary Table 1. Overview of regions captured by the custom CRC panel.

| **Region** | **Chromosome** | **Start position (hg38)** | **Stop position (hg38)** |
| --- | --- | --- | --- |
| **NRAS** | chr1 | 114713777 | 114714014 |
|  | chr1 | 114715992 | 114716239 |
| **TCF7L2** | chr10 | 113141129 | 113141434 |
|  | chr10 | 113152254 | 113152542 |
|  | chr10 | 113157894 | 113158137 |
|  | chr10 | 113165429 | 113165705 |
| **KRAS** | chr12 | 25225504 | 25225843 |
|  | chr12 | 25227184 | 25227467 |
|  | chr12 | 25245199 | 25245483 |
| **TP53** | chr17 | 7670551 | 7670845 |
|  | chr17 | 7673406 | 7673927 |
|  | chr17 | 7674056 | 7674351 |
|  | chr17 | 7674761 | 7675365 |
|  | chr17 | 7675871 | 7676194 |
| **SOX9** | chr17 | 72123467 | 72124262 |
| **SMAD2** | chr18 | 47841718 | 47841958 |
|  | chr18 | 47848378 | 47848698 |
| **SMAD4** | chr18 | 51065384 | 51065673 |
|  | chr18 | 51078209 | 51078516 |
| **PIK3CA** | chr3 | 179198931 | 179199247 |
|  | chr3 | 179203626 | 179203917 |
|  | chr3 | 179218166 | 179218433 |
|  | chr3 | 179234161 | 179234430 |
| **FBXW7** | chr4 | 152324113 | 152324436 |
|  | chr4 | 152325968 | 152326244 |
|  | chr4 | 152326273 | 152326338 |
|  | chr4 | 152328108 | 152328446 |
| **APC** | chr5 | 112780699 | 112781011 |
|  | chr5 | 112792319 | 112792644 |
|  | chr5 | 112801214 | 112801414 |
|  | chr5 | 112815384 | 112815658 |
|  | chr5 | 112819104 | 112819434 |
|  | chr5 | 112826999 | 112827316 |
|  | chr5 | 112827829 | 112828031 |
|  | chr5 | 112828754 | 112829081 |
|  | chr5 | 112834864 | 112835108 |
|  | chr5 | 112837419 | 112840443 |
|  | chr5 | 112840744 | 112840984 |
| **BRAF** | chr7 | 140753212 | 140753464 |
| **FAM123B** | chrX | 64191239 | 64191626 |
|  | chrX | 64191954 | 64192094 |
|  | chrX | 64192139 | 64192353 |
| **Targeted SNP regions** | chr1 | 187641648 | 187641793 |
|  | chr5 | 52897381 | 52897528 |
|  | chr10 | 16504977 | 16505132 |
|  | chr8 | 144439940 | 144440088 |
|  | chr14 | 22052713 | 22052869 |
|  | chr5 | 176529433 | 176529591 |
|  | chr8 | 98021021 | 98021162 |
|  | chr17 | 2364760 | 2364919 |
|  | chr3 | 105533456 | 105533594 |
|  | chr19 | 1488066 | 1488211 |
|  | chr11 | 47835625 | 47835760 |
|  | chr14 | 104940618 | 104940773 |
|  | chr1 | 40738822 | 40738964 |
|  | chr11 | 33043777 | 33043913 |
|  | chr7 | 4860379 | 4860530 |
|  | chr1 | 118101729 | 118101869 |
|  | chrX | 49236990 | 49237130 |
|  | chrY | 10174188 | 10174308 |

### Supplementary section 2: Data generation

For each sample a .fastQ-file containing the paired-end sequencing data is generated. This is mapped to the Hg38 reference genome using BWA-MEM v0.7.17 [4]. The unique Molecular Identifiers (UMI) are then grouped together using the “directional” method in UMI-tools v1.1.1 [5]. Using FilterConsensusReads from fgbio v1.3.0 (<http://fulcrumgenomics.github.io/fgbio/>) the UMI groups are then filtered to a minimum size of three and positions with an error rate above 10% are N-masked. Consensus reads are formed using CallMolecularConsensusReads (fgbio). The consensus reads converted to .fastQ using SamToFastq from picard v2.25.0 [6] and then re-mapped to the reference genome using BWA-MEM. Tags containing UMI specific information are retained using MergeBamAlignment from picard. The read-mapping (.bam) is then filtered to the target region of the panel using samtools v1.13 [7] and reads that are unmapped and not proper-paired or a secondary alignment are removed. Finally, the 5’ and 3’ ends of reads are trimmed by two and one base pairs respectively with ClipBam (fgbio), as these positions have shown to contain a considerable number of errors.

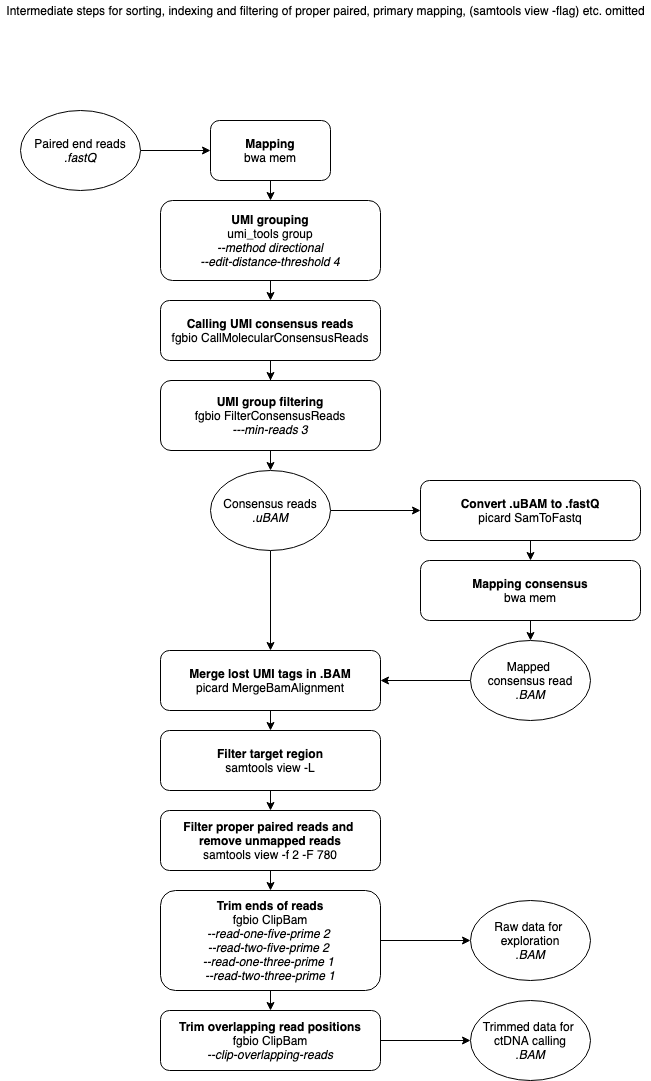

### Supplementary section 3: Explorative analysis of features and error rates

To explore a feature and how it affects the behavior of errors at a read position, the observed distribution of that feature is compared between the set of matches and mismatches. By comparing these we can see some feature values are more prominent in either of the categories, indicating that the feature is useful for explaining the behavior of errors. For a given feature, let $p_{x}$ and $q_{x}$ be the observed frequency of the feature value $x$ in the set of errors and matches respectively. Using an analysis approach akin to naïve Bayes, we can get an estimate of the error rate for that given value of the feature, $\tilde{e}_{x}$, using Bayes’ formula:

$$\tilde{e}_{x}=\frac{p_{x}\cdot e}{p_{x}\cdot e+q_{x}\cdot\left( 1-e \right)}$$

Where $e$ is the observed error rate in the dataset. Using this we can visualize if the estimated error is inflated for some values of the feature, indicating that it could be useful for differentiation between high and low error rate read positions. This can easily be generalized to analyze the interaction between multiple features. However, this analysis approach is naïve with respect to possible correlations with other features since it only considers features explicitly explored in the analysis.

The variance of the error rate estimate can be estimated using the delta method:

$$Var\left( p_{x} \right)\approx\frac{1}{N}p_{x}\left( 1-p_{x} \right), Var\left( q_{x} \right)\approx\frac{1}{M}q_{x}\left( 1-q_{x} \right)$$

$$Var\left( \tilde{e}_{x} \right)\approx\left( \frac{d}{dp_{x}}\tilde{e}_{x} \right)^{2}\cdot Var\left( p_{x} \right)+\left( \frac{d}{dq_{x}}\tilde{e}_{x} \right)^{2}\cdot Var\left( q_{x} \right)$$

Where $N$ and $M$ are the number of observed errors and matches respectively. Assuming that $\tilde{e}_{x}$ is approximately normally distributed, confidence intervals can be calculated using the standard error:

$$SE=\Phi\left( 1-\frac{\alpha}{2} \right)\cdot\sqrt{Var\left( \tilde{e}_{x} \right)}, CI_{\alpha}\left( \tilde{e}_{x} \right)=\left[ \tilde{e}_{x}\pm SE \right]$$

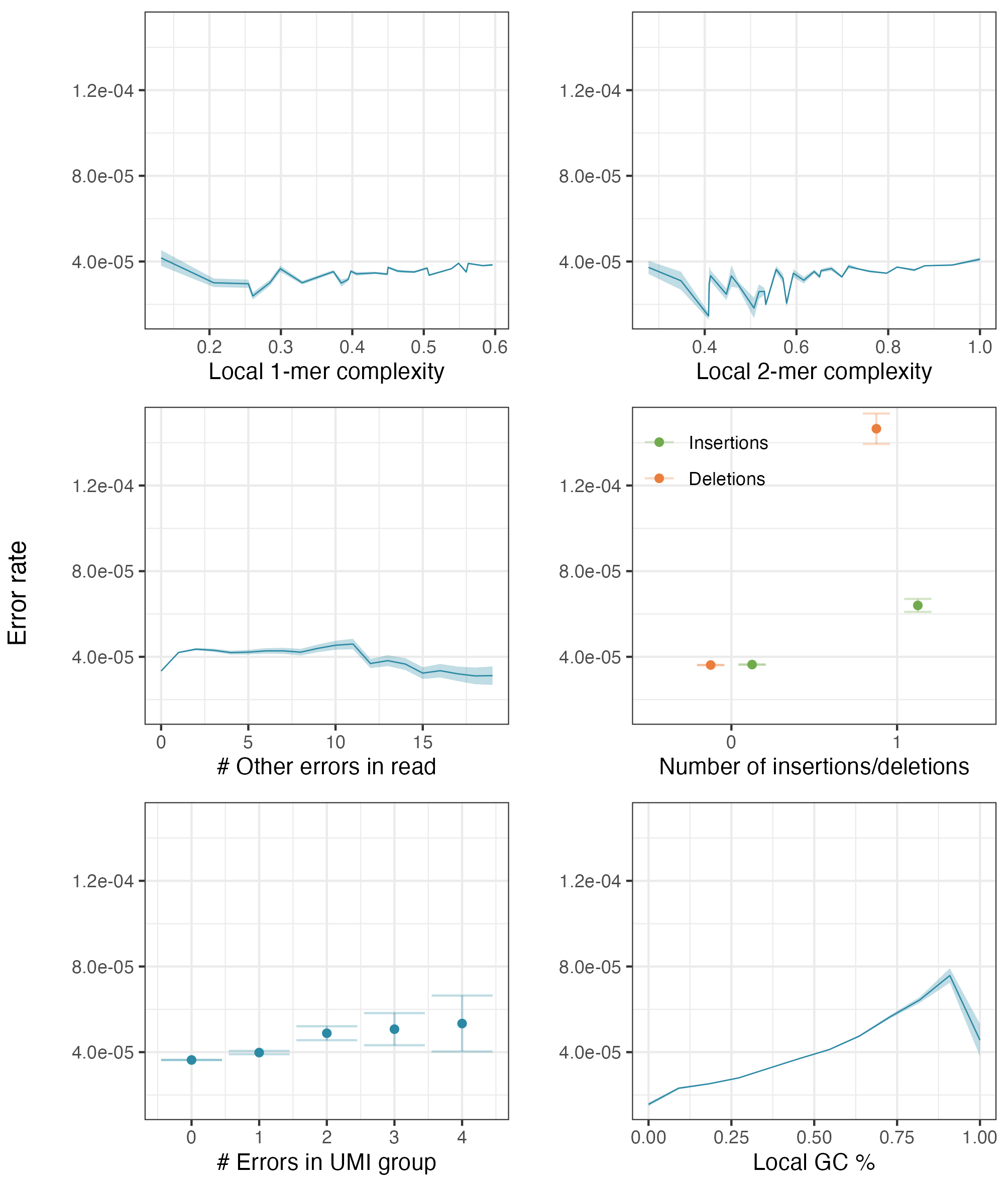

### Supplementary section 4: Feature selection

#### Feature importance

An initial set of features of interest was chosen for the model. Some of these features might not be informative for the task of estimating allelic error rates, and ideally these should be removed from the model to both lower the amount of noise stemming from these features and to limit the complexity of the model. To do this the importance of the individual features was assessed using Leave One Covariate Out (LOCO) as described in [8]. Using this method, the model is trained on a dataset and the predictive performance is measured on a holdout validation dataset. The importance of the $j$’th feature is then estimated as the drop in predictive performance when this process is repeated, but with $j$’th feature removed from the training dataset.

To do this, we start by splitting the data into a training and validation split $D_{Train}$ and $D_{Validation}$ respectively, and train a baseline model, $p^{baseline}$ on the full training data, $D_{Train}$. Similarly, a model, $p^{j}$, is trained on $D_{Train}$, but where the $j$’th feature have been removed. The drop in predictive performance can then be calculated for each individual data point in $D_{Validation}$ as the difference in loss:

$$\Delta_{i}^{j}=l\left( p^{j}\left( x_{i} \right),y_{i} \right)-l\left( p^{baseline}\left( x_{i} \right),y_{i} \right), i\in D_{Validation}$$

Where $l\left( \hat{p},y \right)$ is the categorical cross entropy between the observation $y$ and the predicted probability $\hat{p}$. The feature importance score is then defined as the average difference in loss:

$${FI}^{j}=\frac{1}{\left| D_{Validation} \right|}\sum_{i\in D_{Validation}} \Delta_{i}^{j}$$

Confidence intervals for the feature importance can be created using a gaussian approximation:

$$\left[ {FI}^{j}\pm\Phi^{-1}\left( 1-\frac{\alpha}{2\cdot F} \right)\cdot\frac{s^{j}}{\sqrt{\left| D_{Validation} \right|}} \right], s^{j}=\sqrt{\frac{\sum_{i\in D_{Validation}} \left( FI^{j}-\Delta_{i}^{j} \right)^{2}}{\left| D_{Validation} \right|-1}}$$

, where $F$ is the number of features investigated, used to make a Bonferroni correction of the gaussian quantiles used for the interval.

#### Recursive backwards elimination

If some features contain approximately the same information, removing any one of them would not decrease the performance of the model. This mean that all these features would receive a low features importance score, as they in practice cover for each other’s absence from the model. Thus, only one feature should be eliminated from the model at a time based on the features importance scores. Selecting the final subset of features will therefore be done using a greedy approach, where features are recursively eliminated, in the order of importance as described in the previous section. This idea is similar to what is presented in [9]. After each elimination the average difference in loss is calculated to see if the performance has been affected. Like the calculation of feature importance, we do a 5-fold cross validation at each step to exhaust the information in the dataset. The final model is the one with the fewest features that is not worse than the full model in any folds of the CV. This ensures that the features included in the final model are important in at least one fold, thus only including features that show a large effect on the error rate. For significance testing the Bonferroni used to correct for the number of test (5 folds x 14 features). The full algorithm is as follows:

1. Calculate feature importance for all features using CV.
2. Until no features are left:
   a. Remove the remaining feature with lowest importance.
   b. Train model only using the remaining features.
   c. Calculate the average difference in loss compared to full model
3. Choose the smallest model that is not worse than the full model.

### Supplementary section 5: Down sampling and rescaling

For a neural network to learn the behavior of the errors, this naturally must train on data that has both positive and negative examples. Optimally this training data should simply contain all individual read positions from a set of training sample. However, this immediately present two problems. First, this implies an unpractically large amount of data points. Secondly, the number of non-errors would far exceed the number of errors resulting in a heavily skewed dataset, which is generally undesirable when working with learning-based algorithms. As the number of errors in a dataset is manageable, we can solve both problems by simply down sampling the non-errors in the dataset. However, this drastically biases the estimated class probabilities, as the errors become relatively more prominent in the data. In [10] they explore this problem, and present a procedure for how to re-scale the learned class probabilities using that the down sampling ratio is known. However, this procedure only covers the simple binary case. In the following we present how this procedure is adapted to the current setting, together with a method for obtaining the individual error rates by transforming the binary outcome of this procedure.

To model the down sampling let $S\in\{0,1\}$ represent whether a read position is included in the down sampled dataset (1) or not (0), and let $\beta$ be the probability that a non-error is included:

$$\beta=P\left( S=1|X=R \right)$$

Since the errors are not down sampled, we know that $P\left( S=1|X\neq R \right)=1$. Here $\beta$ can be chosen such that the desired ratio of errors to non-error is obtained. Assuming we want $k$ non-error for every error in the down sampled dataset:

$$\beta=\frac{k\cdot n_{errors}}{T-n_{errors}}$$

, where $T$ is the total number of read positions in the dataset. Now assume $a\neq R$ is an alternative allele and that $e^{R\to a}$ is the corresponding estimated error rate. Since the procedure assumes a binary classification problem, the rates for three possible errors are grouped together. Let $e^{R}$ be the combined error rate:

$$e^{R}=\sum_{a:a\neq R} e^{R\to a}$$

Rewriting the combined error rate conditioning on the down sampling $S$ and using Bayes’ formula yields:

$$e^{R}=P\left( X\neq R|S=1 \right)$$

$$=\frac{P\left( S=1|X\neq R \right)P\left( X\neq R \right)}{P\left( S=1|X\neq R \right)P\left( X\neq R \right)+P\left( S=1|X=R \right)P\left( X=R \right)}$$

$$=\frac{P\left( X\neq R \right)}{P\left( X\neq R \right)+P\left( S=1|X=R \right)P\left( X=R \right)}$$

$$=\frac{\tilde{e}^{R}}{\tilde{e}^{R}+\beta\cdot\left( 1-\tilde{e}^{R} \right)}$$

, where $\tilde{e}^{R}$ is the combined error rate in the original dataset. Using this we get the following relationship between the combined error rates in the original and down sampled dataset:

$$\tilde{e}^{R}=\frac{\beta\cdot e^{R}}{\beta\cdot e^{R}-e^{R}+1}$$

To obtain the individual error rates after correcting for down sampling bias in the training data, we make following novel addition to the procedure. Assume that the proportionality between error rates should stay the same before and after re-scaling, such that:

$$\frac{e^{R\to a}}{e^{R}}=\frac{\tilde{e}^{R\to a}}{\tilde{e}^{R}}, a\neq R$$

Using this together with the individual error rates learned in the down samples data, yields the following individual error rates:

$$\tilde{e}^{R\to R}=1-\tilde{e}^{R}, \tilde{e}^{R\to a}=\tilde{e}^{R}\cdot\frac{e^{R\to a}}{e^{R}}, a\neq R$$

### Supplementary section 6: Estimating tumor fraction

#### Estimating the tumor fraction

In this section we develop the Expectation-Maximization algorithm for estimating the tumor fraction ($f$) and mutation presence probability ($r$) as presented in the model above [11]. To start the analysis let $\left\{ \left( \left\{ \left( x_{ij},y_{ij} \right) \right\}_{j=1}^{N_{i}},z_{i} \right) \right\}_{i=1}^{K}$ be a set of fragments with observed nucleotides, $x_{ij}$, true nucleotides $y_{ij}$ and true mutation status $z_{i}$. It is assumed that the fragments are pairwise independent and that $y_{ij}$ and $z_{i}$ are latent variables since they cannot be observed in practice. This fact will be used later in the E-step of the algorithm. Using this the complete-data log-likelihood function can be written as follows:

$$l\left( f,r | \left\{ \left( \left\{ \left( x_{ij},y_{ij} \right) \right\}_{j=1}^{N_{i}},z_{i} \right) \right\}_{i=1}^{K} \right)=\sum_{i=1}^{K} \log P\left( Z_{i}=z_{i} \right)+\sum_{j=1}^{N_{i}} \log P\left( Y_{ij}=y_{ij}|Z_{i}=z_{i} \right)+\log P\left( X_{ij}=x_{ij}|Y_{ij}=y_{ij} \right)$$

Given this expression the expectation and maximization step of algorithm can developed in the following sections to find maximum likelihood estimates of $\hat{f}$ and $\hat{r}$ in the model above.

##### E-step

In this step the expected value of the complete-data log-likelihood function is calculated. Since $y_{ij}$ and $z_{i}$ are latent variables, their state will be inferred given the assumptions of their distribution in the model and that we have observed data $\left\{ x_{ij} \right\}_{j=1}^{N_{i}}$. Let $f^{t}$ and $r^{t}$ be the current parameter estimates, which are found in the M-step below. The expected complete-data log-likelihood function at step $t$ is:

$$Q^{t}\left( f,r | \left\{ \left( \left\{ \left( x_{ij},y_{ij} \right) \right\}_{j=1}^{N_{i}},z_{i} \right) \right\}_{i=1}^{K} \right)=E_{Y,Z|X}^{t}\left[ l\left( f,r | \left\{ \left( \left\{ \left( x_{ij},y_{ij} \right) \right\}_{j=1}^{N_{i}},z_{i} \right) \right\}_{i=1}^{K} \right) \right]$$

$$=\sum_{i=1}^{K} P_{Y,Z|X}^{t}\left( Z_{i}=0 \right)\cdot\log\left( 1-r \right)+P_{Y,Z|X}^{t}\left( Z_{i}=1 \right)\cdot\log\left( r \right)+$$

$$\quad\quad\sum_{j=1}^{N_{i}} P_{Y,Z|X}^{t}\left( Y_{ij}=R,Z_{i}=0 \right)\cdot$$

$$\quad\quad\quad\quad\left[ 1\left( x_{ij}=R \right)\cdot\log\left( 1-e_{ij}^{R\to M} \right)+1\left( x_{ij}=M \right)\cdot\log\left( e_{ij}^{R\to M} \right) \right]+$$

$$\quad\quad\quad\quad P_{Y,Z|X}^{t}\left( Y_{ij}=R,Z_{i}=1 \right)\cdot$$

$$\quad\quad\quad\quad\left[ 1\left( x_{ij}=R \right)\cdot\log\left( 1-e_{ij}^{R\to M} \right)+1\left( x_{ij}=M \right)\cdot\log\left( e_{ij}^{R\to M} \right)+\log\left( 1-\frac{f}{2} \right) \right]+$$

$${\quad\quad\quad\quad P}_{Y,Z|X}^{t}\left( Y_{ij}=M,Z_{i}=1 \right)\cdot$$

$$\quad\quad\quad\quad\left[ 1\left( x_{ij}=M \right)\cdot\log\left( 1-e_{ij}^{M\to R} \right)+1\left( x_{ij}=R \right)\cdot\log\left( e_{ij}^{M\to R} \right)+\log\left( \frac{f}{2} \right) \right]$$

The crucial part of the E-step is to calculate the probabilities of the states of the latent variables in this expression. Since it generally holds that:

$$P_{Y,Z|X}^{t}\left( Y_{ij}=y_{ij},Z_{i}=z_{i} \right)=P_{Y,Z|X}^{t}\left( Y_{ij}=y_{ij} | Z_{i}=z_{i} \right)\cdot P_{Y,Z|X}^{t}\left( Z_{i}=z_{i} \right)$$

We can focus on calculating the two factors on right site separately in the following. We start by exploring the second factor:

$$P_{Y,Z|X}^{t}\left( Z_{i}=0 \right)=\frac{P^{t}\left( \left\{ x_{ij} \right\}_{j=1}^{N_{i}} | Z_{i}=0 \right)\cdot P^{t}\left( Z_{i}=0 \right)}{P^{t}\left( \left\{ x_{ij} \right\}_{j=1}^{N_{i}} \right)}$$

For now let $\frac{1}{c}=P^{t}\left( \left\{ x_{ij} \right\}_{j=1}^{N} \right)$. We can then rewrite the expression above:

$$P_{Y,Z|X}^{t}\left( Z_{i}=0 \right)=c\cdot\left( 1-r^{t} \right)\prod_{j=1}^{N} P^{t}\left( x_{ij} | Z_{i}=0 \right)$$

$$=c\cdot\left( 1-r^{t} \right)\prod_{j:x_{ij}=R} \left( 1-e_{ij}^{R\to M} \right)\cdot\prod_{j:x_{ij}=M} e_{ij}^{R\to M}$$

Similarly, we get:

$$P_{Y,Z|X}^{t}\left( Z_{i}=1 \right)=c\cdot r^{t}\cdot\prod_{j:x_{ij}=R} \left[ \left( 1-e_{ij}^{R\to M} \right)\cdot\left( 1-\frac{f^{t}}{2} \right)+e_{ij}^{M\to R}\cdot\frac{f^{t}}{2} \right]\cdot\prod_{j:x_{ij}=M} \left[ e_{ij}^{R\to M}\cdot\left( 1-\frac{f^{t}}{2} \right)+\left( 1-e_{ij}^{M\to R} \right)\cdot\frac{f^{t}}{2} \right]$$

Note that $c$ can found using the fact that $P_{Y,Z|X}^{t}\left( Z_{i}=0 \right)+P_{Y,Z|X}^{t}\left( Z_{i}=1 \right)=1$.

We then turn our focus to the conditional probability of the true fragment state:

$$P_{Y,Z|X}^{t}\left( Y_{ij}=y_{ij} | Z_{i}=z_{i} \right)=P^{t}\left( Y_{ij}=y_{ij} | X_{ij}=x_{ij},Z_{i}=z_{i} \right)$$

$$=\frac{P^{t}\left( X_{ij}=x_{ij} | Y_{ij}=y_{ij} \right)\cdot P^{t}\left( Y_{ij}=y_{ij} | Z_{i}=z_{i} \right)}{P^{t}\left( X_{ij}=x_{ij} | Z_{i}=z_{i} \right)}$$

And from this we get:

$$P_{Y,Z|X}^{t}\left( Y_{ij}=R | Z_{i}=0 \right)=\left\{ \begin{matrix} 1 & x_{ij}=R \\ 1 & x_{ij}=M \end{matrix} \right.$$

$$P_{Y,Z|X}^{t}\left( Y_{ij}=M | Z_{i}=0 \right)=\left\{ \begin{matrix} 0 & x_{ij}=R \\ 0 & x_{ij}=M \end{matrix} \right.$$

$$P_{Y,Z|X}^{t}\left( Y_{ij}=R | Z_{i}=1 \right)=\left\{ \begin{matrix} \frac{\left( 1-e_{ij}^{R\to M} \right)\cdot\left( 1-\frac{f^{t}}{2} \right)}{\left( 1-e_{ij}^{R\to M} \right)\cdot\left( 1-\frac{f^{t}}{2} \right)+e_{ij}^{M\to R}\cdot\frac{f^{t}}{2}} & x_{ij}=R \\ \frac{e_{ij}^{R\to M}\cdot\left( 1-\frac{f^{t}}{2} \right)}{e_{ij}^{R\to M}\cdot\left( 1-\frac{f^{t}}{2} \right)+\left( 1-e_{ij}^{M\to R} \right)\cdot\frac{f^{t}}{2}} & x_{ij}=M \end{matrix} \right.$$

$$P_{Y,Z|X}^{t}\left( Y_{ij}=M | Z_{i}=1 \right)=\left\{ \begin{matrix} \frac{e_{ij}^{M\to R}\cdot\frac{f^{t}}{2}}{\left( 1-e_{ij}^{R\to M} \right)\cdot\left( 1-\frac{f^{t}}{2} \right)+e_{ij}^{M\to R}\cdot\frac{f^{t}}{2}} & x_{ij}=R \\ \frac{\left( 1-e_{ij}^{M\to R} \right)\cdot\frac{f^{t}}{2}}{e_{ij}^{R\to M}\cdot\left( 1-\frac{f^{t}}{2} \right)+\left( 1-e_{ij}^{M\to R} \right)\cdot\frac{f^{t}}{2}} & x_{ij}=M \end{matrix} \right.$$

##### M-step

In this step the function $Q^{t}$ is maximized to get updated parameter estimates $f^{t+1}$ and $r^{t+1}$, using the probabilities of the states of the latent variables found in the E-step above. To maximize the function with respect to $f$ the derivative is calculated:

$$\frac{d}{df}Q^{t}\left( f,r | \left\{ \left( \left\{ \left( x_{ij},y_{ij} \right) \right\}_{j=1}^{N_{i}},z_{i} \right) \right\}_{i=1}^{K} \right)=\frac{-1}{2-f}\sum_{i=1}^{K} \sum_{j=1}^{N_{i}} P_{Y,Z|X}^{t}\left( Y_{ij}=R,Z_{i}=1 \right)+\frac{1}{f}\cdot\sum_{i=1}^{K} \sum_{j=1}^{N_{i}} P_{Y,Z|X}^{t}\left( Y_{ij}=M,Z_{i}=1 \right)=0$$

Yielding the updating function:

$$f^{t+1}=2\cdot\frac{\sum_{i=1}^{K} \sum_{j=1}^{N_{i}} P_{Y,Z|X}^{t}\left( Y_{ij}=M,Z_{i}=1 \right)}{\sum_{i=1}^{K} \sum_{j=1}^{N_{i}} P_{Y,Z|X}^{t}\left( Y_{ij}=M,Z_{i}=1 \right)+\sum_{i=1}^{K} \sum_{j=1}^{N_{i}} P_{Y,Z|X}^{t}\left( Y_{ij}=R,Z_{i}=1 \right)}$$

$$=2\cdot\frac{\sum_{i=1}^{K} P_{Y,Z|X}^{t}\left( Z_{i}=1 \right)\sum_{j=1}^{N_{i}} P_{Y,Z|X}^{t}\left( Y_{ij}=M | Z_{i}=1 \right)}{\sum_{i=1}^{K} P_{Y,Z|X}^{t}\left( Z_{i}=1 \right)\cdot N_{i}}$$

Note that this can be interpreted as double the observed frequency of fragments with the mutated allele in mutated positions, which is consistent with the way $f$ was introduced in the model as the tumor fraction. Doing the same with respect to $r$ we get:

$${\frac{d}{dr}Q}^{t}\left( f,r | \left\{ \left( \left\{ \left( x_{ij},y_{ij} \right) \right\}_{j=1}^{N_{i}},z_{i} \right) \right\}_{i=1}^{K} \right)=\frac{-1}{1-r}\cdot\sum_{i=1}^{K} P_{Y,Z|X}^{t}\left( Z_{i}=0 \right)+\frac{1}{r}\cdot\sum_{i=1}^{K} P_{Y,Z|X}^{t}\left( Z_{i}=1 \right)=0$$

Yields the following updating function:

$$r^{t+1}=\frac{\sum_{i=1}^{K} P_{Y,Z|X}^{t}\left( Z_{i}=1 \right)}{\sum_{i=1}^{K} P_{Y,Z|X}^{t}\left( Z_{i}=1 \right)+\sum_{i=1}^{K} P_{Y,Z|X}^{t}\left( Z_{i}=0 \right)}$$

$$=\frac{\sum_{i=1}^{K} P_{Y,Z|X}^{t}\left( Z_{i}=1 \right)}{K}$$

This has the nice interpretation of the fraction of sites in the catalogue that are mutated, which is how $r$ was introduced in the model.

##### Algorithm initialization

To initialize the algorithm, the following heuristic is used. First, find minimum and maximum values for each of the parameters $f$ and $r$, then do a coarse grid search for a good pair of starting values. For $f$ the maximum value is chosen as twice the maximum observed frequency:

$$f_{max}=\max_{i} \left( 2\cdot\frac{\sum_{j=1}^{N_{i}} x_{ij}}{N_{i}} \right)$$

For $r$ the minimum is chosen as the case where only one mutation is present, and the maximum value is chosen as the ratio of positions with observed signal:

$$r_{min}=\frac{1}{K}, r_{max}=\frac{\sum_{i=1}^{K} 1\left( \sum_{j=1}^{N_{i}} x_{ij}>0 \right)}{K}$$

The grid search is then done by evaluating the log-likelihood function at point where, $f=f_{max}\cdot{10}^{k}$ for different values of $k<0$ and $r$ is chosen by splitting the interval $(r_{min},r_{max})$ uniformly.

#### Confidence intervals

Furthermore, confidence intervals for $f$ and $r$ is constructed by looking at significance testing marginally for each parameter. In general, the confidence interval for a parameter $\theta$ becomes the region around the MLE, $\hat{\theta}$, for which the significance test of the null hypothesis $H_{0}:\theta=\theta_{0}$ is not rejected for a given $\alpha$-level. For this model this corresponds to calculating the following intervals:

$$CI_{f}^{\alpha}=\left\{ f | Q\left( f,\hat{r} \right)\leq F_{\chi^{2}\left( 1 \right)}^{-1}\left( 1-\alpha\right) \right\}, CI_{r}^{\alpha}=\left\{ r | Q\left( \hat{f},r \right)\leq F_{\chi^{2}\left( 1 \right)}^{-1}\left( 1-\alpha\right) \right\},$$

$$Q\left( f,r \right):=-2\log\frac{L\left( \hat{f},\hat{r} | \left\{ \left( x_{ij} \right) \right\}_{j=1}^{N_{i}} \right)}{L\left( f,r | \left\{ \left( x_{ij} \right) \right\}_{j=1}^{N_{i}} \right)}$$
